# Characterization of *Azotobacter vinelandii* and Kits for Its Synthetic Biology Applications

**DOI:** 10.1101/188482

**Authors:** King Pong Leung, Jacky Fong Chuen Loo, Leo Chi U Seak, Tung Faat Lai, Kevin Yuk Lap Yip, Siu Kai Kong, Ting Fung Chan, King Ming Chan

## Abstract

*Azotobacter vinelandii*, a Gram-negative aerobic bacterium with an intracellular anaerobic environment that maintains the oxygen-sensitive enzymatic cascades for nitrogen fixation, could be used to express oxygen-sensitive proteins. However, little is known about the properties of *A. vinelandii* for synthetic biology applications. We therefore first characterized and optimized the conditions for growing and screening BioBrick constructs in *A. vinelandii* in the presence of 2 antibiotics, ampicillin and chloramphenicol, and then developed two sets of BioBricks for regulated protein expression. The first kit used T7 RNA polymerase, whose expression is under the control of a nitrogen-repressible *nifH* promoter. The commonly used T7-dependent system in *Escherichia coli* can then be used in *A. vinelandii*. Because its intracellular anaerobic environment is favorable for processes such as magnetosome biogenesis, we attempted to migrate the biogenesis machineries from the magnetotactic bacterium *Magnetospirillum gryphiswaldense* to *A. vinelandii*. During this undertaking, another insertion kit construct was developed to allow protein conjugation onto magnetosomes. The kit consists of *mamC*, a gene encoding a transmembrane protein on magnetosomes, and multiple restriction sites downstream of *mamC* for fusing a gene of interest. This insertion kit allows the attachment of any desired protein onto the magnetosome membrane by fusing with the mamC protein. We demonstrated the function of this kit by fusing mamC to a GFP nanobody. This kit will facilitate the conjugation of any target protein onto magnetosomes for downstream applications in the future.

**Financial Disclosure:** We received sponsorship from the 2012–15 Teaching Development Grants Triennium, Faculty of Engineering and Biochemistry Program, School of Life Sciences, The Chinese University of Hong Kong. The funders had no role in the study design, data collection and analysis, decision to publish, or preparation of the manuscript.

**Competing Interests:** The authors declare that no competing interests exist.

**Ethics Statement:** N/A.

**Data Availability:** All data are fully available without restriction.

## Introduction

*Azotobacter vinelandii*, a Gram-negative nitrogen-fixing aerobic bacterium, has been shown to create an intracellular anaerobic environment to protect its oxygen-sensitive nitrogenase complexes [1]. The mechanism operates by consuming intracellular oxygen with enhanced respiratory enzymatic activity [2] and preventing molecular oxygen diffusion by extracellular alginate [3]. These features make the bacterium a potential host for the production and characterization of oxygen-sensitive proteins or organelles. Transformation of large size DNA is possible through stable genome integration [4], and common antibiotics such as ampicillin and chloramphenicol can be used in *A. vinelandii* for selection after transformation. However, no previous studies have reported the use of this bacterium in synthetic biology applications. Our idea is to use *A. vinelandii* to express recombinant proteins under anaerobic conditions. We report some basic characterizations and applications of synthetic constructs of *A. vinelandii*.

Recombinant protein expression in *E. coli* often adopts the IPTG-inducible T7 expression system in which T7 RNA polymerase is produced under IPTG induction, which in turn transcribes gene(s) that is under the control of a T7 promoter. Because there is no endogenous T7 RNA polymerase in *A. vinelandii*, we designed a T7 protein expression construct using a *nifH* promoter whereby the expression of the downstream T7 RNA polymerase is repressed by ammonium (NH_4_^+^). Nitrogenase in *A. vinelandii* is most highly expressed when the nitrogen source is depleted and is inhibited when a fixed nitrogen source (i.e., NH_4_^+^) is present [5] because of the nitrogen-dependent activation of the *nif* operon regulated by nifL and nifA. Because the *nifH* promoter is under the regulation of these two proteins, putting T7 RNA polymerase under its control should create a protein expression system that is suppressed by the addition of NH_4_^+^ and derepressed when NH_4_^+^ is removed. The constructs were registered as BBa_K1314013 and BBa_K1314014 in the Registry of Standard Biological Parts (http://parts.registry.org). They bear a strong ribosomal binding site (RBS) (BBa_B0034) and a weak RBS (BBa_B0031) for translation regulation, respectively.

To demonstrate that anaerobic processes such as magnetosome formation can be run in *A. vinelandii*, an aerobe having an intracellular anaerobic environment, we attempted to produce magnetosomes by migrating necessary machineries from magnetotactic bacteria. These bacteria form intracellular membrane-bound organelles called magnetosomes. The diameter of each vesicle encapsulating a magnetic crystal composed of magnetite (Fe_3_O_4_) or greigite (Fe_3_S_4_) is about 35–120 nm [6]. Fusion of the target protein with a magnetosome may provide a range of applications, such as co-immunoprecipitation assays. *Magnetospirillum gryphiswaldense* is one of the well-studied magnetotactic bacteria. However, it is microaerophilic in nature, and microaerobic/anaerobic growing conditions are critical for magnetosome formation [1,7]. We sought to migrate the magnetosome biogenesis machinery of *M. gryphiswaldense* to *A. vinelandii*, a potential aerobic host providing an intracellular anaerobic environment for easy manipulation with standard laboratory equipment. Therefore, in the course of introducing magnetosome biogenesis operons into *A. vinelandii*, we developed an insertion kit construct for protein expression on magnetosomes that consists of the *mamC* gene encoding a transmembrane protein on magnetosomes [8] and multiple restriction sites (MCS) downstream of *mamC* for fusing a gene of interest.

## Methods

### Strains and Growth Conditions

For molecular cloning, chemically competent *E. coli* DH5α (Life Technologies) was used, which was cultured in LB medium at 37°C at 250 rpm. *A. vinelandii* strain DJ was kindly provided by Dr. Dennis R. Dean. *A. vinelandii* was cultured in Burk’s medium (B medium: 0.2% KH2PO4, 0.8% K2HPO4, 20% sucrose, 0.2% MgSO_4_·7H_2_O, 0.09% CaCl_2_·2H_2_O, 0.005% FeSO_4_·7H_2_O, 0.1 nM Na_2_MoO_4_·2H_2_O and 1.6% agar) [10] or with the addition of 13 mM NH4OAc (BN medium) at 30°C at 300 rpm. The minimal inhibitory concentration of antibiotics (ampicillin or chloramphenicol) was determined by measuring the optical density at 600 nm (OD600) of the bacterial culture after 12-h exposure to various drug concentrations as indicated.

### Confocal Microscopy

Bacterial culture (10 μl) was immobilized by mixing it with 10 μl of 1% low-melting-point agarose. Confocal fluorescence images were captured with a Leica TCS SP8 confocal laser scanning system with a 60× objective lens (Leica Microsystems, Germany). Yellow fluorescent protein (YFP) was excited with a 488-nm laser source, and the fluorescence signal was detected at the range of 495–545 nm.

### Plasmid Construction

Phusion^®^ DNA polymerase for PCR and all restriction endonucleases were purchased from New England Biolabs. The PCR target sizes were validated by gel electrophoresis. DNA sequencing of target bands was provided by the BGI DNA sequencing platform (Hong Kong).

#### (A) Nitrogen-repressible T7 Expression System

The *nifH* promoter and *nifK* homologous sequences were amplified from *A. vinelandii* genomic DNA with the primer sequences listed in **Table 1**. By double digestion and ligation according to iGEM standard RFC10, we put BBa_K1314000 (*nifH* promoter) upstream of BBa_K1314005 (T7 polymerase with a strong RBS) and BBa_K1314006 (T7 polymerase with a weak RBS). BBa_K1314012 (*nifK* homologous sequence) and BBa_K1314002 (150-bp random sequence) were then respectively added to both ends of the DNA sequence. Also, BBa_R0040 (tetR_pro_), BBa_B0034 (RBS), BBa_E1010 (red fluorescent protein [RFP]) and BBa_B0015 (double terminator) were added downstream of the construct as a reporter.

**Table 1.**
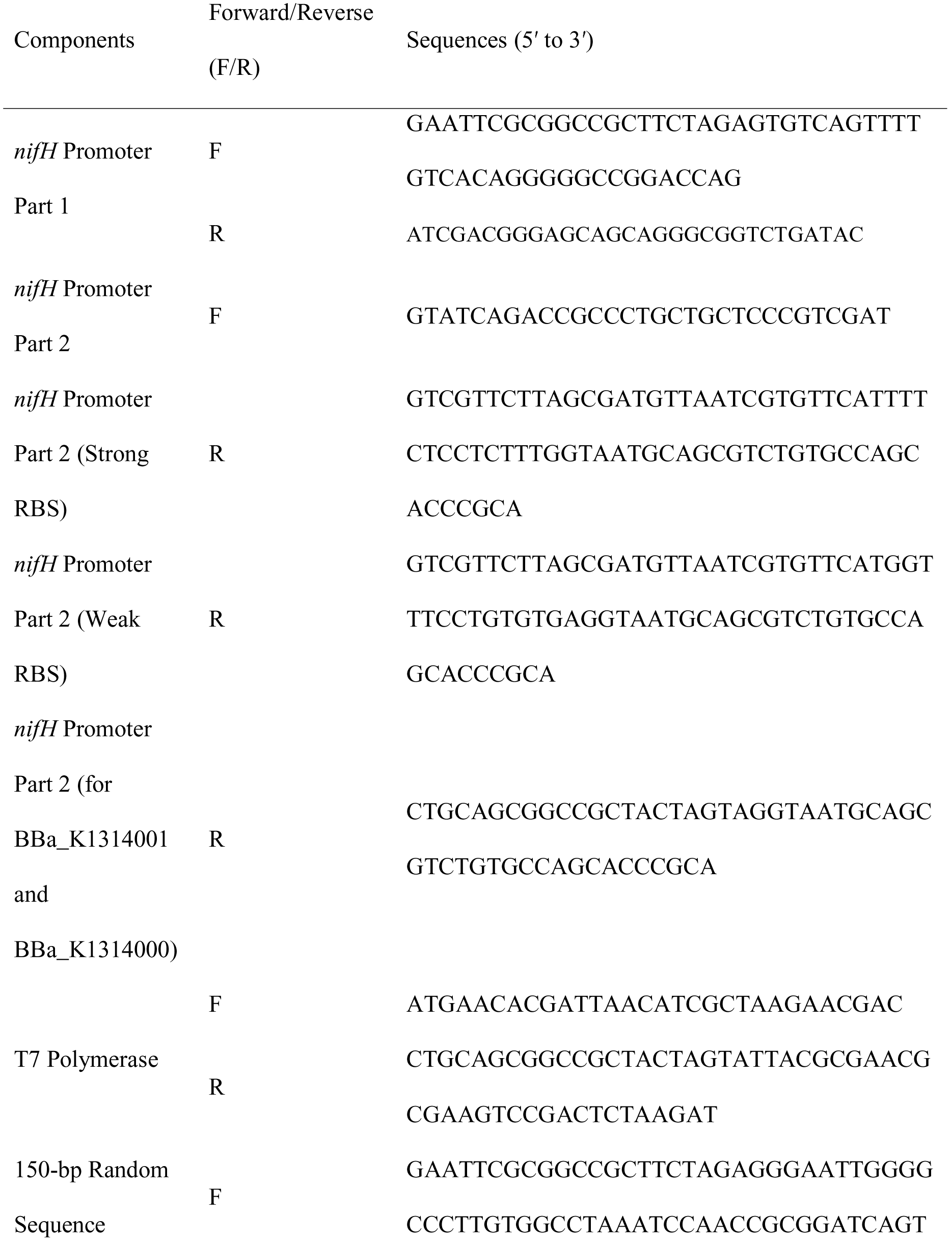
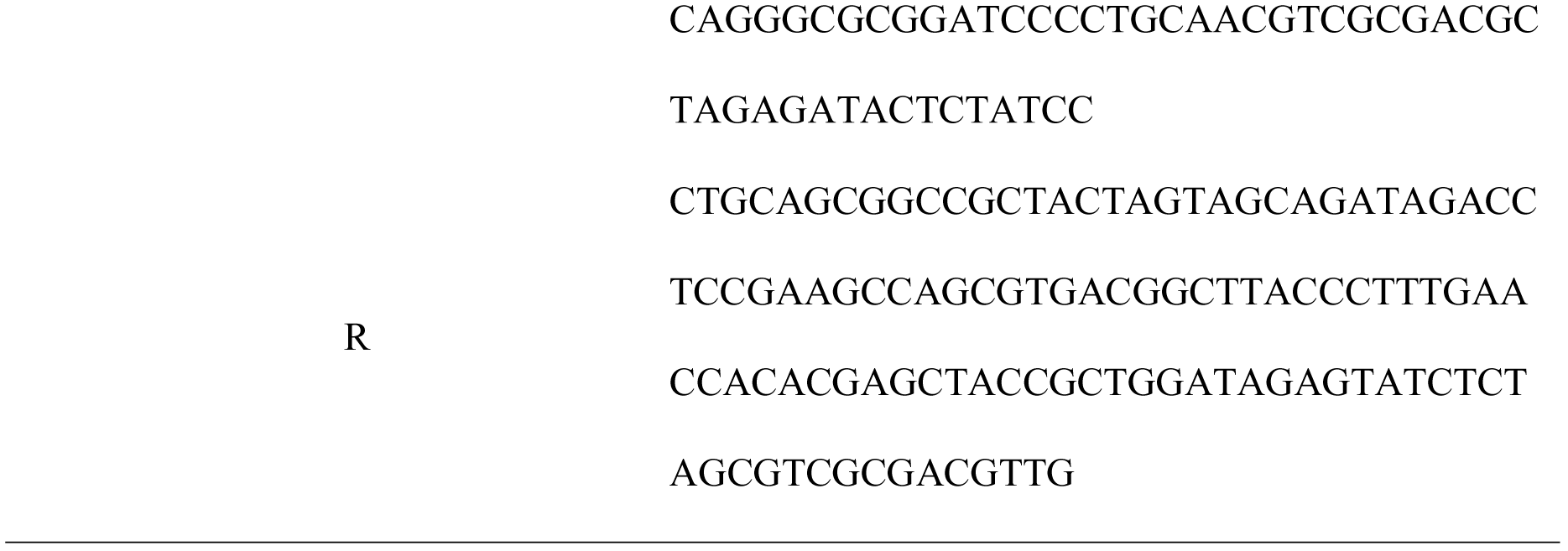
Primers for Constructing the Nitrogen-repressible T7 Expression System.

#### (B) Insertion Kit for Expressing Protein Expression on the Magnetosome Membrane

An overview of the cloning strategy is outlined in **Figure 4A**. Four fragments were made via PCR, namely the pSB1C3 plasmid backbone, BBa_J13002 (consisting of the constitutive promoter tetR_pro_ BBa_R0040 and the RBS), BBa_B0015 (double terminator) and mamC, using the primers listed in **Table 2**. By using overlapping PCR of these four fragments, one linear fragment was produced. The PCR reactions were subjected to a three-step cyclic program (98°C for 1 min, followed by 35 cycles of 98°C for 10 s, 72°C for 30 s with a 0.5°C decrease per cycle, and 72°C for 10 min). PstI sites were added at both ends of the linear fragment, allowing a circular plasmid (BBa_K1648004) to be produced by ligating the PstI-digested fragment.

**Figure 4.**
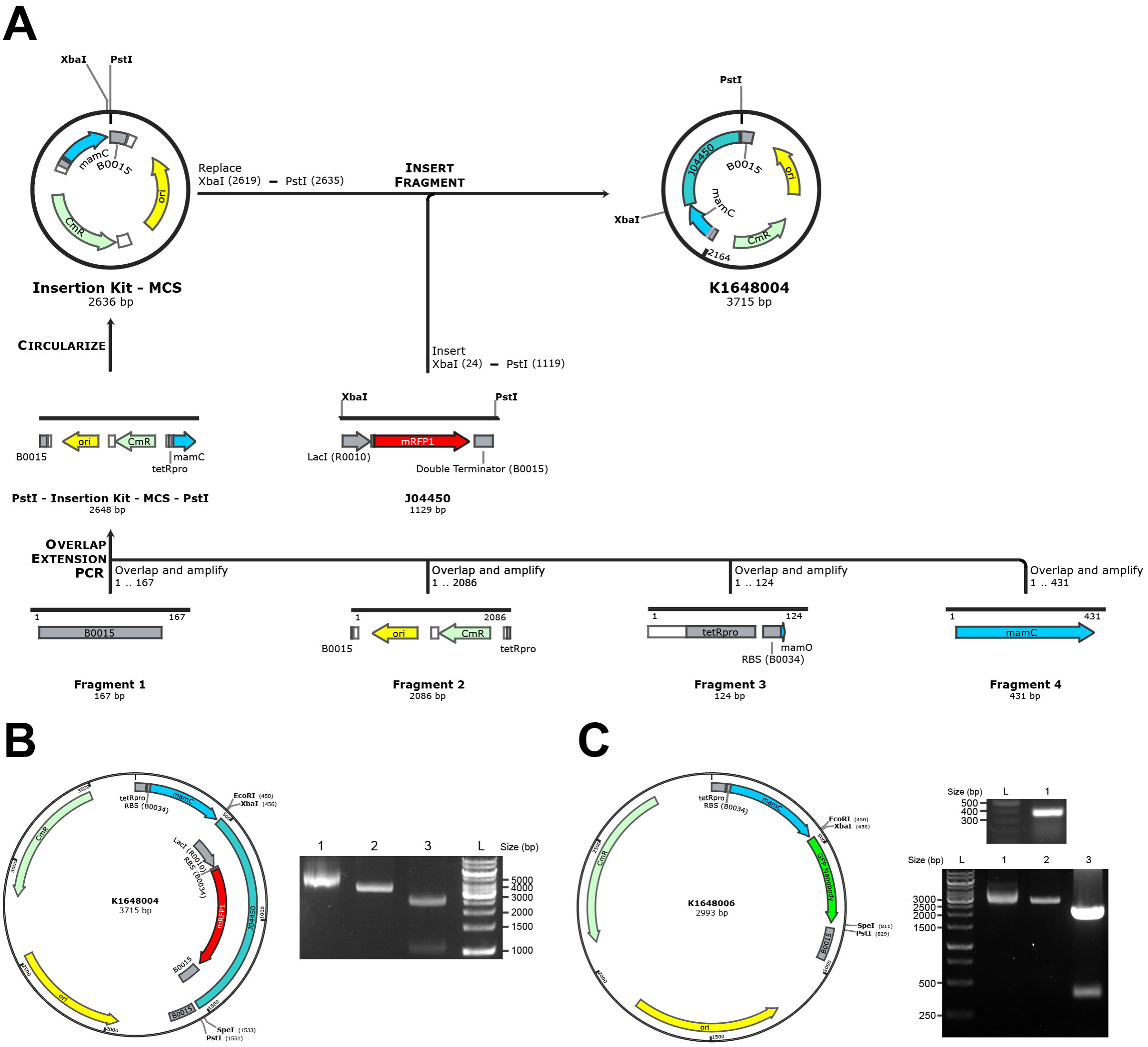
Cloning strategy for construction of an insertion kit and construct maps of the kit (BBa_K1648004) with GFP nanobody (BBa_K1648006). **(A)** Flow diagram showing the cloning strategy for the insertion kit. **(B)** Insertion kit for expressing a protein of interest on magnetosomes (BBa_K1648004). Left panel: Construct map of BBa_K1648004. Right panel: Gel electrophoresis of the insertion kit for fusing a protein of interest on the magnetosome membrane (BBa_K1648004) without digestion (Lane 1), with single digestion at the XbaI site (Lane 2) and with double-digestion cut at the XbaI and PstI sites (Lane 3). **(C)** Insertion kit with GFP nanobody fused with mamC (BBa_K1648006). Left panel: Construct map of BBa_K1648006. Lower right panel: Gel electrophoresis of (Upper) PCR product of GFP nanobody with size of 400 bp; (Lower) BBa_K1648006 without digestion (Lane 1), with single digestion at the XbaI site (Lane 2) and with double-digestion cut at the XbaI and PstI sites (Lane 3). L represents DNA ladder.

**Table 2.**
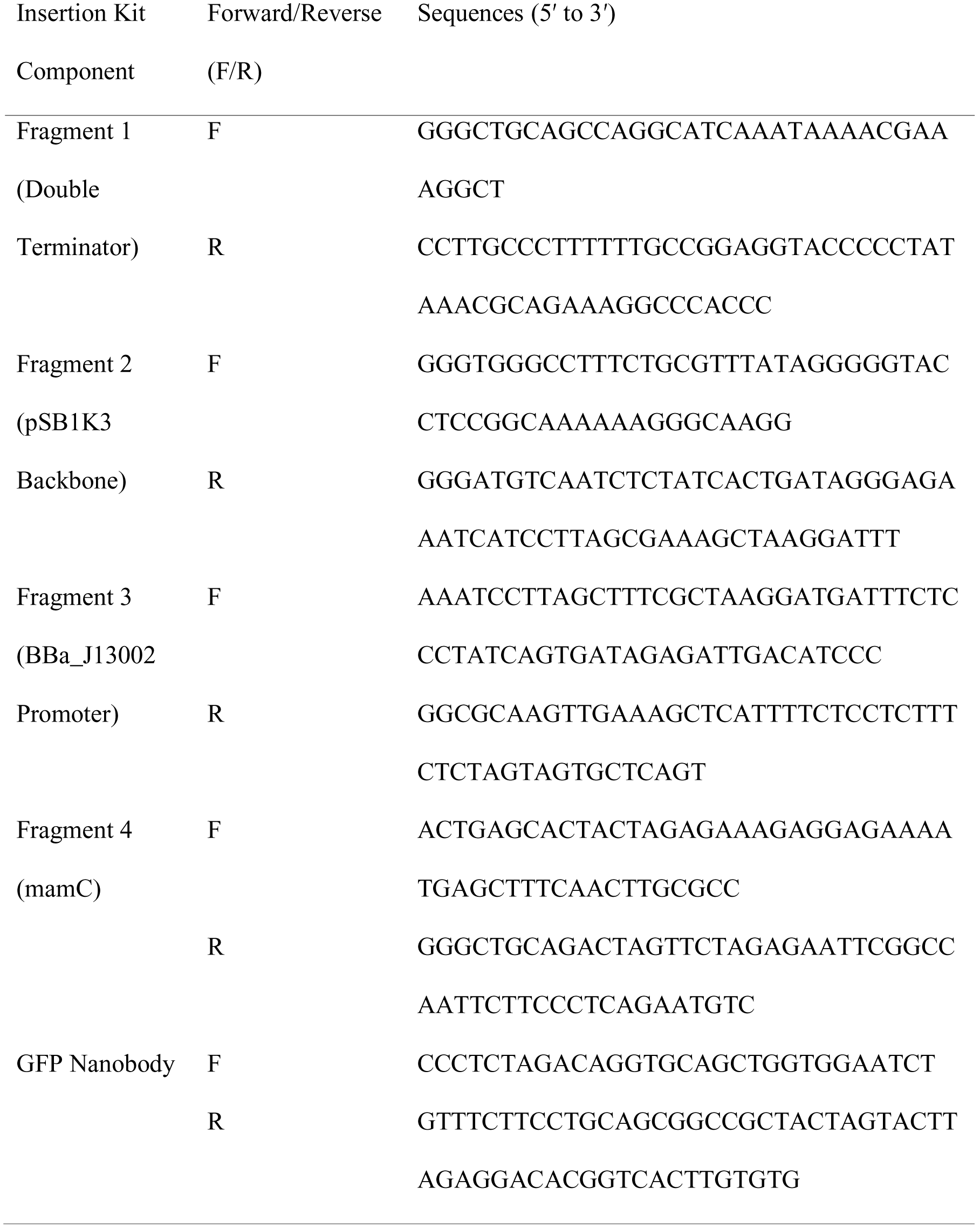
Primers for Constructing the Insertion Kit for Protein Expression on the Magnetosome Membrane.

### *Natural Transformation of* A. vinelandii

Transformation of *A. vinelandii* was performed with a protocol from Dr. Dennis R. Dean (personal communication) with modifications. Briefly, *A. vinelandii* cells were streaked onto BN-Mo plates (0.2% KH_2_PO_4_, 0.8% K_2_HPO_4_, 20% sucrose, 0.2% MgSO_4_·7H_2_O, 0.09% CaCl_2_·2H_2_O, 0.005% FeSO_4_·7H_2_O, 13 mM NH_4_OAc and 1.6% agar) and were grown for 2 days in a 30°C incubator. A loop of cells was added to a 125-ml sterile flask containing the medium composed of 0.2% KH_2_PO_4_, 0.8% K_2_HPO_4_, 20% sucrose, 0.2% MgSO_4_·7H_2_O, 0.09% CaCl_2_·2H_2_O and 65 μl of 5 M NH_4_OAc. The flask was set on a bench for 5 min, followed by vortexing for 10 s. The culture was then grown for 18–20 h at 30°C and 170 rpm. Resultant competent cells (200 μl) were mixed with 200 μl of 1× MOPS transformation buffer (20 mM MOPS and 20 mM MgSO_4_·7H_2_O) and 5 μl plasmid DNA. The mixture (300 μl) was incubated at room temperature for 15–20 min, and 2.7 ml of B medium was then added into the mixture. The mixture containing the cells was shaken overnight and then spread on a B medium plate with 1.6% agar and appropriate antibiotics.

## Results

### *Duration and Minimum Antibiotics Dosage Required for* A. vinelandii *Transformant Selection*

We first determined the optimal duration for growing *A. vinelandii* liquid culture. The growth curve of wild-type *A. vinelandii* grown at 30°C at 300 rpm is shown in **Figure 1A**. Surprisingly, the log-phase growth of *A. vinelandii* lasted beyond 24 h and up to 36 h (data not shown). We also found that it was optimal to subculture every 3 days to maintain exponential growth (data not shown).

**Figure 1.**
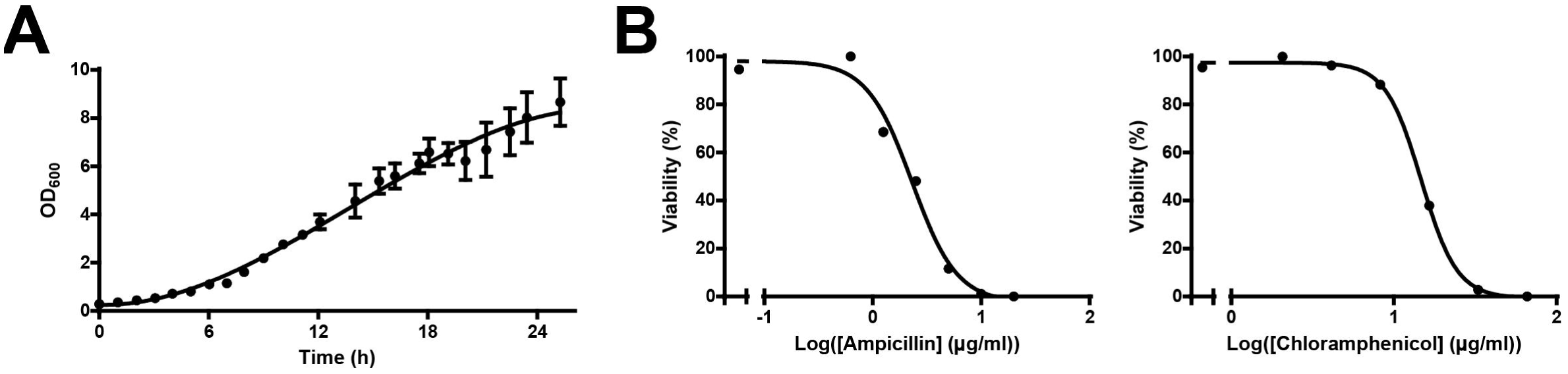
Conditions for growing and selecting transformants by antibiotics in *A. vinelandii*. **(A)** Growth curve of *A. vinelandii*. Bacterial culture was grown at 30°C at 300 rpm. OD_600_ was monitored at the indicated time intervals. **(B)** Growth inhibition of *A. vinelandii* under different concentrations of antibiotics (ampicillin or chloramphenicol) as indicated. OD_600_ was measured after 12 h of treatment.

To determine the minimal antibiotics dosage that could eliminate non-transformants, we applied a range of concentrations of two commonly used antibiotics, ampicillin and chloramphenicol, to wild-type *A. vinelandii*. We observed 100% growth inhibition at 10 μg/ml of ampicillin and at 66 μg/ml of chloramphenicol (**Figure 1B**). Therefore, the concentrations used for screening transformants using these antibiotics should be beyond the minimal dosage required. In the following experiments, we added ampicillin or chloramphenicol in BN medium at the concentrations of 25 μg/ml and 100 μg/ml, respectively.

### *Anaerobic Intracellular Environment in* A. vinelandii

It has been suggested that the oxygen-labile nitrogenase enzyme complex essential for nitrogen fixation is protected by two mechanisms in *A. vinelandii* [1]. To confirm whether the intracellular environment is free of molecular oxygen, i.e. anaerobic, we used YFP as a probe. YFP is derived from GFP and requires oxygen for chromophore maturation [11]. Therefore, YFP only fluoresces when oxygen is present. We expressed YFP residing in cytosol and INP-YFP (ice nucleation protein; BBa_K1423005) localized on the outer membrane in *A. vinelandii*, with *E. coli* serving as a control. YFP was exposed to the extracellular medium when fused to INP; thus, it fluoresced when exposed to the molecular oxygen present in the medium for both *A. vinelandii* and *E. coli*, as shown in **Figure 2**. However, although cytosolic YFP fluoresced in *E. coli*, indicating the presence of oxygen, its counterpart in *A. vinelandii* did not. This result indicated that *A. vinelandii* has little or no molecular oxygen present in its internal environment.

**Figure 2.**
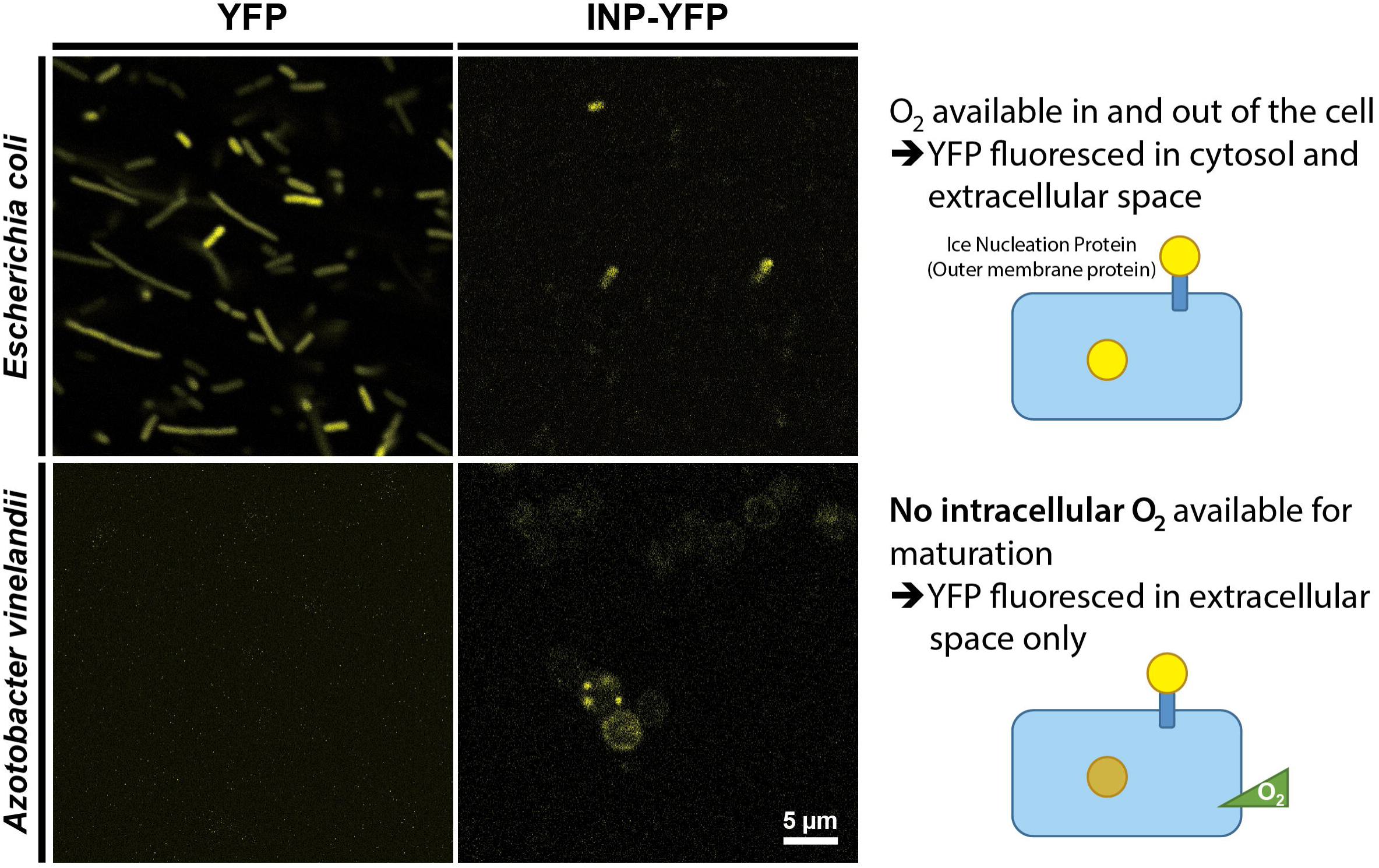
*A. vinelandii* has an intracellular anaerobic environment. YFP was used as a probe to detect molecular oxygen. Both cytosolic YFP and INP-YFP (ice nucleation protein) localized on the outer membrane were expressed in *A. vinelandii* and *E. coli*. YFP was exposed to an extracellular medium when fused to INP, and thus it can fluoresce by exposure to molecular oxygen present in the medium for both *A. vinelandii* and *E. coli*.

### *Two Protein Expression Kits for* A. vinelandii

#### (A) Nitrogen-repressible T7 Expression System for *A. vinelandii*

We constructed a nitrogen-repressible expression system by putting T7 RNA polymerase under the control of the NH4^+^-regulated *nifH* promoter (**Figure 3A**). BBa_K1314013 in the pSB1A3 backbone was linearized with XhoI to facilitate homologous recombination of the *nifH*_pro_::T7 RNA polymerase cascade into the *nifHDK* operon of the *A. vinelandii* genome. Such recombination is achieved with homologous sequences, the *nifH* promoter and a sequence 150 bp from the 3**’** end of the *nifK* gene. An RFP-expressing cascade was added to indicate successful random integration into the *A. vinelandii* genome. pSB1A3 plasmid backbone contains ampicillin resistance gene for selection. Because the *nifHDK* operon encodes the nitrogenase structural genes [12], no active nitrogenase could be made for nitrogen fixation, a process that could lead to suppression of the nitrogen-regulated T7 expression cascade. A 150-bp random sequence, which produced no significant hit by BLASTN against the *A. vinelandii* genomic sequence, was included for subsequent introduction of the T7_pro_::protein-of-interest cascade by homologous recombination. Linearized BBa_K1314011 contains the random sequence and part of the RFP reporter as homologous sequences on both ends for replacement of the original RFP expression cascade in the genome. With a design similar to BBa_K1314011, any T7 expression system used in *E. coli* can be used in *A. vinelandii* as well.

**Figure 3.**
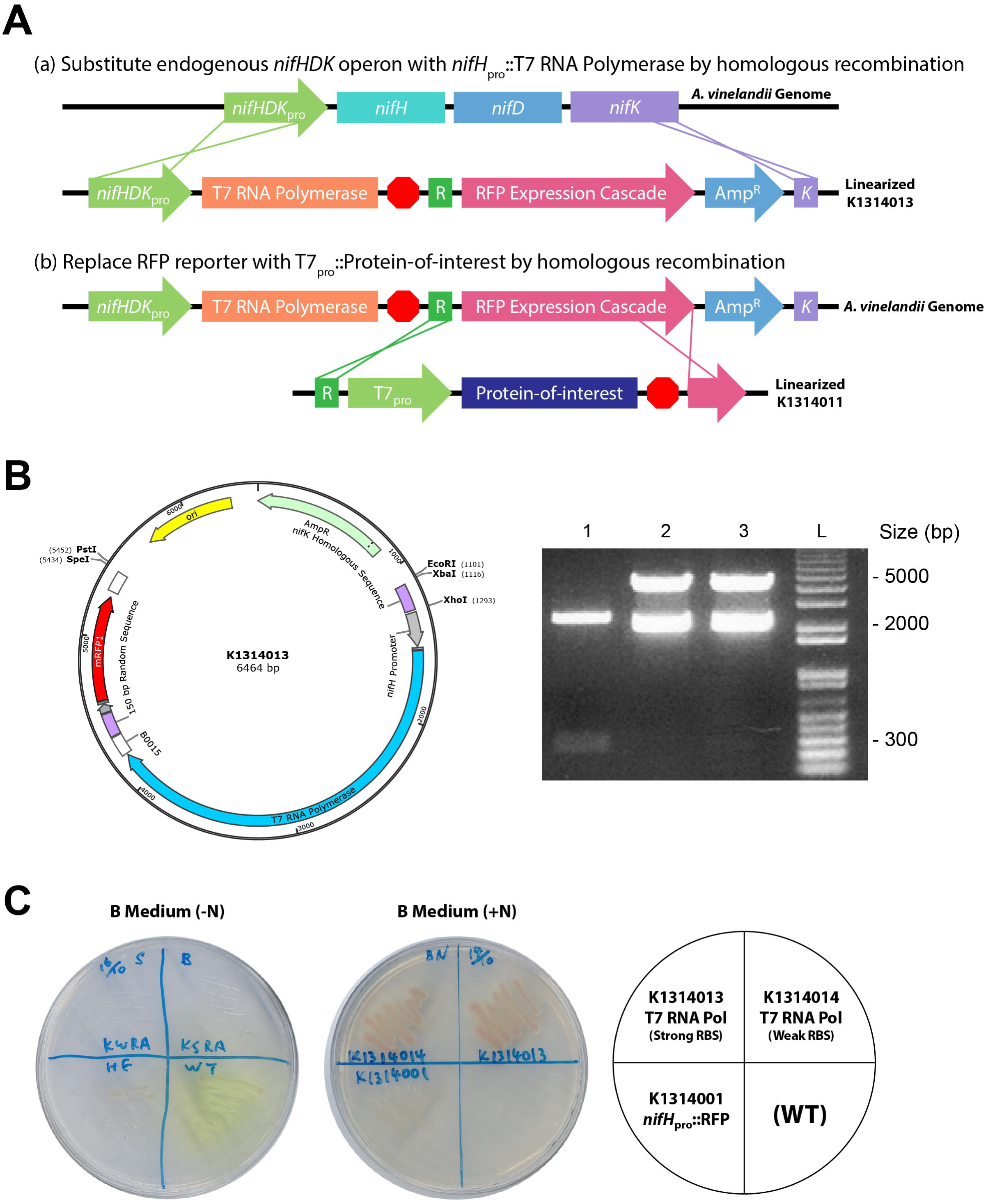
Nitrogen-repressible T7 protein expression system in *A. vinelandii*. **(A)** Schematic diagram showing the design of the T7 protein expression system. Red octagon: double terminator; Green “R”: 150-bp random sequence for homologous recombination; Purple “K”: 150 bp from the 3**’** end of the *nifK* gene for homologous recombination. **(B)** *nifH*_pro_::T7 RNA polymerase cascade for homologous recombination (BBa_K1314013). Left panel: Construct map of BBa_K1314013. XhoI was introduced for linearization. Right panel: Gel electrophoresis of EcoRI-PstI double-digested products of BBa_K1314000 *(nifH* promoter only, Lane 1), BBa_K1314013 (Lane 2) and BBa_K1314014 (Lane 3). L represents DNA ladder. **(C)** Growth of *A. vinelandii* stably integrated with BBa_K1314001 (*nifH*_pro_::RFP), BBa_K1314013 and BBa_K1314014 with or without nitrogen (NH_4_^+^).

The construct map of BBa_K1314013 is shown in **Figure 3B**. BBa_K1314013 harbors a strong RBS, BBa_B0034, whereas another construct, BBa_K1314014, was also made with a weak RBS, BBa_B0031. The plasmids BBa_K1314000 (which contains the *nifH* promoter of about 300 bp only), BBa_K1314013 and BBa_K1314014 were verified by double digestion with EcoRI and PstI. Bands of about 5 kb were observed in both BBa_K1314013 and BBa_K1314014 as expected.

We characterized the growth of BBa_K1314001 (*nifH*_pro_::RFP), BBa_K1314013 and BBa_K1314014 stably integrated in *A. vinelandii* with or without nitrogen (NH_4_^+^). Under nitrogen depletion, wild-type and BBa_K1314001 could grow, and BBa_K1314001 was able to express RFP (**Figure 3C**). This result indicated that the *nifH* promoter allows gene expression under a nitrogen condition. Because the *nifHDK* operon was replaced via homologous recombination with BBa_K1314013 or BBa_K1314014, these two strains cannot grow in the absence of nitrogen. When NH_4_^+^ was present, BBa_K1314001 grew well, but no RFP was expressed, indicating the suppression of the *nifH* promoter under a nitrogen-rich condition. Meanwhile, BBa_K1314013 and BBa_K1314014 grew well with RFP expressed, which suggested successful integration of the T7 RNA polymerase cascades with the RFP reporter through homologous recombination.

#### (B) Insertion Kit for Expressing Protein Expression on the Magnetosome Membrane

We used a cloning strategy of our designed insertion kit construct as shown in **Figure 4A**. It consists of the *mamC* gene from the magnetosome island of the *M. gryphiswaldense* genome, a constitutive promoter tetR_pro_ with strong RBS (BBa_J13002) in front of the MCS, and a double terminator (BBa_B0015) after the MCS. Because the target protein will be fused to the C-terminus of mamC by inserting it into the MCS, the stop codon of mamC was removed. The resultant plasmid is the insertion kit construct registered in the Registry of Standard Biological Parts as BBa_K1648004. The construct map of BBa_K1648004 is shown in **Figure 4B**. The plasmid was verified by PCR and gel electrophoresis with the expected size of 2770 bp, where the PCR band lies in parallel with the ladder between 2500 and 3000 bp. To demonstrate whether a target protein to be expressed could be inserted into the kit, we used GFP nanobody (BBa_K1648005) as an example. BBa_K1648005 was inserted into the MCS of the insertion kit (BBa_K1648004), resulting in the BioBrick expressing the mamC-GFP nanobody (BBa_K1648006). Validation of the BioBrick is shown in **Figure 4C**. The plasmid was verified by double digestion with XbaI and PstI, which gives a 400-bp DNA product as expected.

## Discussion

Given that *A. vinelandii* produces an intracellular anaerobic environment that protects oxygen-sensitive enzymatic cascades and yet can itself be grown aerobically, we believe that this feature could be used in synthetic biology to construct systems that require anaerobic conditions but at the same time allow users to manipulate the cultures easily with standard laboratory equipment. To this end, we attempted to migrate magnetosome biogenesis components from *M. gryphiswaldense*, a biological process that requires little or no oxygen. During the attempt, we developed two kits for protein expression in *A. vinelandii*.

The nitrogen-repressible T7 protein expression system could allow the expression of oxygen-sensitive proteins in the anaerobic intracellular environment inside *A. vinelandii*, a fast-growing aerobe. The *nifHDK* operon, the operon encoding nitrogenase structural genes [12], was intentionally substituted by the T7 RNA polymerase cascade. As nitrogenase is essential for providing a nitrogen source when nitrogen is depleted, knockout of *nifHDK* could prevent the T7 system from being suppressed. Although it seems paradoxical to express protein in a nitrogen-depleted condition, the protein expression can be controlled by titrating the amount of NH_4_^+^ in the growth medium, which can keep the *nifH* promoter functional and allow protein synthesis at the same time. Because the T7 protein expression system is commonly used, this allows easy migration of existing expression cascades, such as BBa_K1497017 (naringenin-producing operon) and BBa_K1465304 (alcohol dehydrogenase), into *A. vinelandii*, either by random genome integration or homologous recombination.

Meanwhile, for the magnetosome protein expression kit, by choosing a GFP nanobody as the protein to be expressed onto magnetosomes, we could capture GFP for easy characterization by detecting its fluorescence. By fusing the GFP nanobody, which has a high binding affinity toward GFP (dissociation constant = 1.4 nM [13]), onto the magnetosome membrane, we could easily extract GFP or GFP-tagged protein by magnetic force. The insertion kit worked as expected with the GFP nanobody inserted. By replacing BBa_J04450 with the desired gene, the protein of interest could be expressed on the magnetosome after transformation of this part into the *A. vinelandii* genome via random integration. After plasmid linearization by restriction endonuclease, the protein could be stably and easily integrated into the *A. vinelandii* genome by random insertion [9]. Therefore, other synthetic biology researchers working with magnetosomes will now be able to use our simple insertion kit to conveniently express various proteins on magnetosomes. Other applications such as protein extraction/purification, which is a series of processes intended to isolate one or a few proteins of interest from a protein mixture, may also be achieved using our insertion kit. Among a myriad of extraction/purification techniques, affinity ligand techniques currently represent the most powerful tool available for downstream processing in terms of their selectivity and recovery. Magnetic separation is usually relatively gentle on target proteins or peptides during extraction/purification, and thus, damage to large protein complexes by traditional column chromatography techniques can be avoided [14].

To conclude, the use of *A. vinelandii* provides a novel method of genetic engineering. It eases the work of engineering a protein or organelle that requires an anaerobic condition. Ultimately, there are numerous possible uses of *A. vinelandii*, and a thorough exploration of the industries that can benefit from its use is warranted.

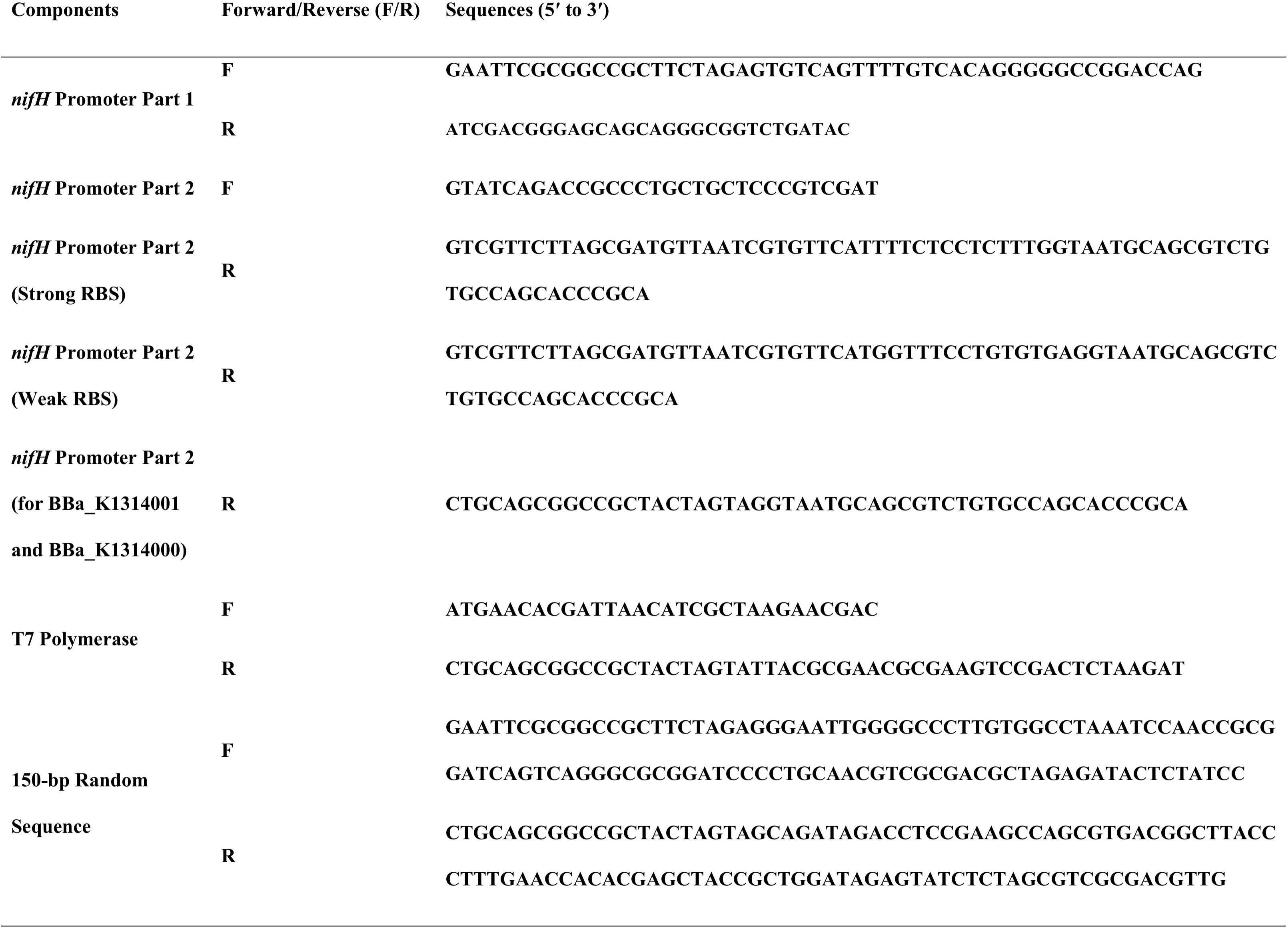
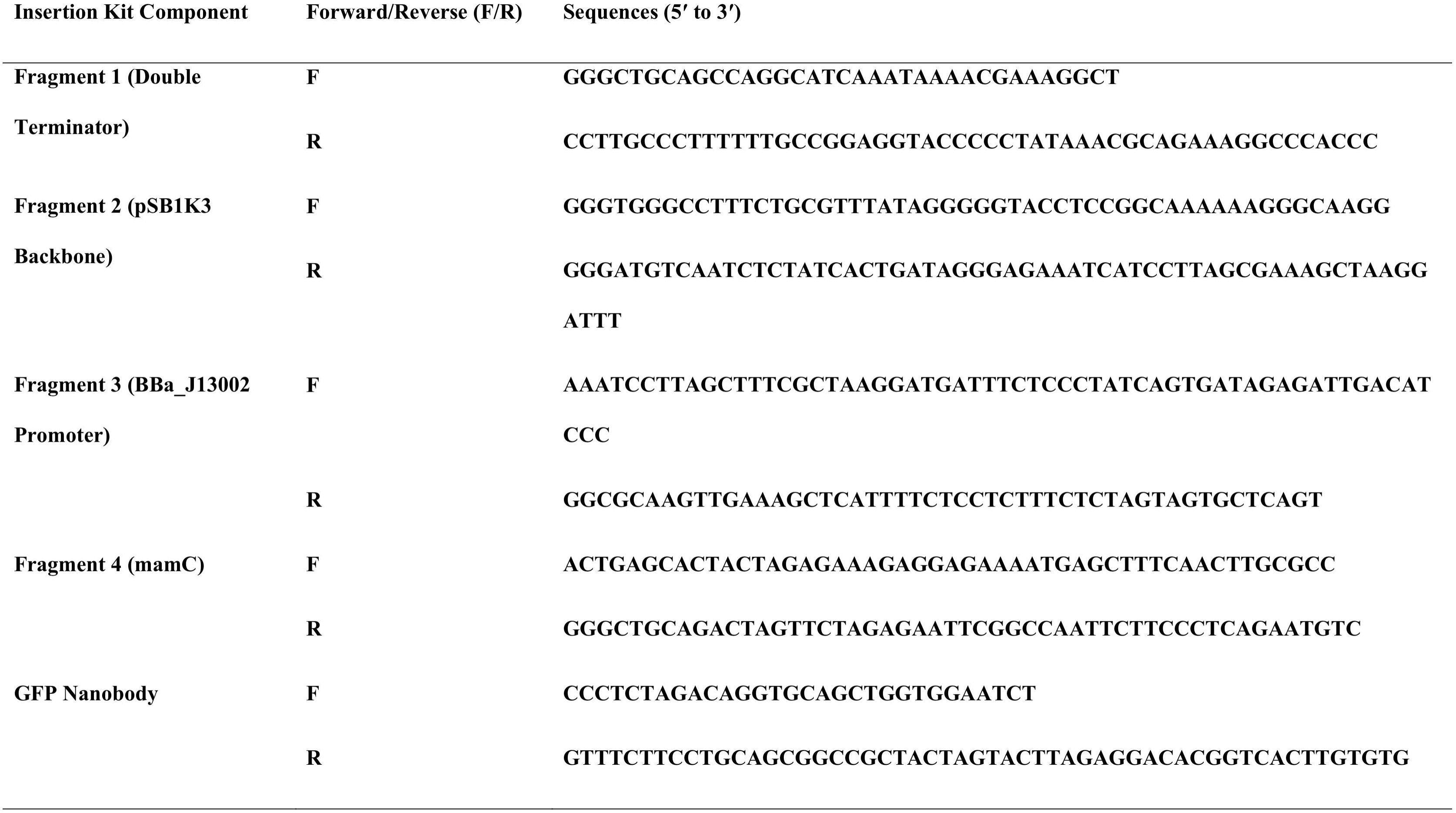

